# Systemic silencing and DNA methylation of a host reporter gene induced by a beneficial fungal root endophyte

**DOI:** 10.1101/2022.06.19.496700

**Authors:** Athanasios Dalakouras, Afrodite Katsaouni, Marianna Avramidou, Elena Dadami, Olga Tsiouri, Sotirios Vasileiadis, Athanasios Makris, Maria Eleni Georgopoulou, Kalliope K. Papadopoulou

## Abstract

A growing body of evidence suggests that RNA interference (RNAi) plays a pivotal role in the communication between plants and pathogenic fungi, where a bi-directional cross-kingdom RNAi is established to the advantage of either the host or the pathogen. Similar mechanisms acting during plant association with non-pathogenic symbiotic microorganisms have been elusive to this date. Here, we report on an RNAi-based mechanism of communication between a beneficial fungal endophyte, *Fusarium solani* strain K (FsK) and its host plants. This soil-borne endophyte that confers resistance and/or tolerance to biotic and abiotic stress in tomato and, as shown in this study, promotes plant growth in *Nicotiana benthamiana*, is restricted to the root system in both host plants. We first showed that the fungus has a functional core RNAi machinery; double stranded RNAs (dsRNAs) are processed into short interfering RNAs (siRNAs) of predominantly 21-nt in size, which lead to the degradation of homologous mRNAs. Importantly, by using an RNAi sensor system, we demonstrated that root colonization of *N. benthamiana* by FsK led to the induction of systemic silencing and DNA methylation of a host reporter gene.. These data reflect a more general but so far unrecognized mechanism wherein root endophytes systemically translocate RNAi signals to the aboveground tissues of their hosts to modulate gene expression during symbiosis, which may be translated to the beneficial phenotypes.

**Highlight:** A root-restricted, beneficial fungal endophyte can induce systemic silencing and epigenetic modifications to its host plant.

## Introduction

RNA interference (RNAi) is a conserved eukaryotic gene regulatory mechanism that is triggered by small RNAs (sRNAs) of approximately 20-25 nucleotides (nt) (Baulcombe, 2004; Hung and Slotkin, 2021). Notwithstanding the diversity of RNAi pathways and the plethora of sRNA classes, there are essentially two types of sRNAs, the small interfering RNAs (siRNAs) and the microRNAs (miRNAs) (Borges and Martienssen, 2015; Vaucheret, 2006). In general, Dicer and Dicer-like (DCL) endonucleases cleave double stranded RNAs (dsRNAs) and stem loop hairpin RNAs (hpRNAs) into 20-25-nt siRNAs and miRNAs, respectively (Paturi and Deshmukh, 2021). The occurring double stranded sRNA is then unzipped in an ATP-dependent reaction so that only one of its two strands will eventually be loaded onto an Argonaute (AGO) protein (Iwakawa and Tomari, 2022; Vaucheret, 2008). Then, the AGO-loaded sRNA scans the cytoplasm for complementary mRNA transcripts to cleave them or inhibit their translation (Brodersen *et al*., 2008; Hamilton and Baulcombe, 1999). At least in plants, AGO-loaded sRNAs may also be transported in the nucleus, where they are involved in RNA-directed DNA methylation (RdDM) of cognate sequences (Wassenegger and Dalakouras, 2021; Wassenegger *et al*., 1994). Moreover, in plants, nematodes and some fungi, the presence of RNA-dependent RNA polymerases (RDRs) contributes to the generation of dsRNAs from single stranded transcripts, in a process termed transitivity (de Felippes and Waterhouse, 2020; Sakurai *et al*., 2021).

Fungal RNAi, initially described as ‘quelling’ in *Neurospora crassa* (Romano and Macino, 1992), has essentially a two-fold role. One the one hand, siRNAs generated from (usually RDR-transcribed) dsRNA precursors are involved in genome defense and maintenance of genome integrity as well as fighting against transposons, viruses and transgenes (Lax *et al*., 2020; Torres-Martinez and Ruiz-Vazquez, 2017). On the other hand, miRNAs (also called miRNA-like, milRNAs), generated by Pol III-transcribed primary miRNA transcripts, fine-tune gene expression during vegetative and sexual development besides responding to various kinds of stresses (Li *et al*., 2010; Torres-Martinez and Ruiz-Vazquez, 2017). A growing body of recent evidence suggests that, in addition to the aforementioned roles, RNAi also has a pivotal role in the communication of fungi with their hosts. Indeed, the pathogen *Botrytis cinerea* delivers sRNAs in Arabidopsis and tomato that target members of the mitogen-activated protein kinases (MAPKs) that function in plant immunity (Weiberg *et al*., 2013). In reverse, plants fight back; Arabidopsis and tomato deliver sRNAs in *B. cinerea* targeting the fungal DCL1 and DCL2, to attenuate fungal pathogenicity and growth (Wang *et al*., 2016). Likewise, *Fusarium graminearum* translocates sRNAs to target defence genes in *Hordeum vulgare* and *Brachypodium distachyon* (Werner *et al*., 2021), whereas cotton plants, in response to infection with the vascular pathogen *Verticillium dahliae*, export miR159 and miR166 to silence fungal isotrichodermin C-15 hydroxylase and Ca(2+)-dependent cysteine protease, respectively, both of which are essential for fungal virulence (Zhang *et al*., 2016). However, the role of such cross-kingdom RNAi processes in mutualistic interactions remains poorly understood.

*Fusarium solani* strain K (FsK) is an endophytic, non-pathogenic strain, initially isolated from the roots of tomato plants (Kavroulakis *et al*., 2007) but other plant species serve as hosts, including legumes (Skiada *et al*., 2019). FsK has been shown to protect the host against root and foliar pathogens (Kavroulakis *et al*., 2007), spider mites (Pappas *et al*., 2018), zoophytophagous insects (Garantonakis *et al*., 2018) and to alleviate drought stress (Kavroulakis *et al*., 2018). The beneficial activity of FsK presupposes an intact ethylene signaling pathway, suggesting that the fungus can induce systemic responses to the plant (Kavroulakis *et al*., 2007). However, the exact molecular details governing this symbiosis remain largely elusive. In this study, we characterized the core RNAi machinery of FsK and provide evidence that the endophyte translocates RNAi signals to its host plant to modulate expression and induce epigenetic modification of a host reporter gene.

## Materials and Methods

### Isolation of fungal conidia and inoculation

FsK was routinely cultured for 4 days in potato dextrose broth (PDB) (26 °C, 160 rpm). Following removal of mycelium fragments by sieving through sterile cheesecloth, conidia were recovered from the filtrate by centrifugation at 6,500 rpm, counted using a haemocytometer and suspended in an appropriate volume of 0.85% NaCl to achieve the desired inoculum concentration. Approximately 100 conidia were used to inoculate *N.benthamiana* plants at cotyledon stage.

### Fungal RNA isolation

FsK was routinely cultured for 4 days in potato dextrose broth (PDB) (26 °C, 160 rpm). From the occuring mycelium total RNA was isolated with TRIzol™ Reagent (www.thermofisher.com) to be subsequently used in RT-qPCR reactions. For small RNA sequencing, the enriched for small RNAs fraction was isolation from the mycelium using mirVana™ miRNA Isolation Kit (www.thermofisher.com) according to the manufacturer’s instructions.

### Reverse transcriptase quantitative polymerase chain reaction (RT-qPCR)

DNaseI-treated (www.thermofisher.com) RNA isolated from mycelium was quantified with by Qubit Fluorometric Quantification (www.thermofisher.com). The DNA-free RNA (10 ng) was then subjected to RT-qPCR using the the Luna® Universal Probe One-Step RT-qPCR Kit (www.neb.com) according to the manufacturer’s instructions. Essentially, the total volume of the reaction was reduced to 10μl and the cycling parameters consisted of incubation at 55°C for 10 min for reverse transcription, 95°C for 1 min followed by 39 cycles of 95°C for 10 sec and 60°C for 30 sec. Analysis was carried out using the geometric mean of FsK ITS and Tef-1a transcripts (Skiada *et al*., 2019). For Tef-1a (120 bp amplicon), the primers 5’-TCG AAC TTC CAG AGG GCA AT-3’ and 5’-CCA ACA ATA GGA AGC CGC TG-3’ were used. For ITS (108 bp amplicon), the primers 5’-TAG GGT AGC TGG GTC TGA CT-3’ and 5’-ACC AAG TCT AAC CCG CCT AC-3’ were used. For GFP (133 bp amplicon), the primers 5’-TCC CAG CAG CTG TTA CAA AC-3’ and 5’-AAT ACT CCA ATT GGC GAT GG-3’ were used. The relative expression of GFP gene was calculated from two to three technical replicates for every sample as described in the corresponding figure legend. Data were analyzed using the Student’s two-tailed homoscedastic t-test.

### Plant and fungal DNA isolation

Genomic DNA from plant and fungal tissue was isolated with DNeasy Plant Pro (/www.qiagen.com) according to the manufacturer’s instructions.

### Phylogenetic analysis

The analyzed sequences were aligned with MUSCLE v3.7 (Edgar, 2004), and informative sites were selected with Gblocks v0.91b (Talavera and Castresana, 2007). The aligned selected sites were tested with the Prottest v3.2 software (Darriba et al., 2011) using the Akaike information criterion (AIC) values for optimal residue substitution model matrix selection. The LG (Le and Gascuel, 2008) residue substitution model matrix scored best for all proteins sets. The PhyML v3.0 algorithm (Guindon and Gascuel, 2003) using the LG model and bootstrap testing with 100 replicates was used for obtaining the best maximum likelihood tree.

### Quantification of fungal colonization by qPCR

To estimate fungal abundance within plant tissues, absolute quantification of *F. solani* ITS gene was performed as previously described (Skiada *et al*., 2019).

### Generation of constructs

For the generation of pCS-GFP, a PCR was performed using as template genomic DNA from *N. benthamiana* line 16C (Voinnet and Baulcombe, 1997) and the primers 5-GGT TAA CAA AGA ATG CTA ACC-3 and 5-CGA GCT CGG CAA TTC CCG ATC-3 and the occuring 2017 bp amplicon was cleaved with HpaI/SacI and ligated to a similarly cleaved pSilent-1, generating the pSilent-GFP. Next, pSilent-GFP was cleaved with PsiI/SacI and the 6663 bp fragment was ligated into the 7866 bp fragment retrieved upon ZraI/SacI cleavage of pCambia1300, generating the pCS-mGFP. For the generation of pCS-hpGF+GFP, a first PCR was performed using as template genomic DNA from *N. benthamiana* line 16C and the primers 5-acg tct cga gAT GAA GAC TAA TCT TTT TCT C-3 and 5-ACG TAA GCT TCT CTT GAA GAA GTC GTG CCG C-3 and the occuring 340 bp amplicon was cleaved with XhoI/HindIII and ligated to a similarly cleaved pSilent-1 vector, generating the pSilent-GF. A second PCR was performed using as template genomic DNA from *N. benthamiana* line 16C and the primers 5-acg tgg tac cAT GAA GAC TAA TCT TTT TCT C-3 and 5-ACG TAG ATC TCT CTT GAA GAA GTC GTG CCG C-3 and the occuring 340 bp amplicon was cleaved with KpnI/BglII and ligated to a similarly cleaved pSilent-GF vector, generating the pSilent-hpGF. Next, the 1937 bp fragment emerging upon HpaI/SacI cleavage of the pSilent-GFP was ligated into a similarly cleaved pSilent-hpGF, generating the pSilent-hpGF+GFP. Finally, pSilent-hpGF+GFP was cleaved with PsiI/SacI and the 7277 bp fragment was ligated into the 7866 bp fragment retrieved upon ZraI/SacI cleavage of pCambia1300, generating the pCS-hpGF+GFP.

### Agrobacterium-mediated fungal transformation

The binary vectors pCS-GFP and pCS-hpGF+GFP were used to transform *Agrobacterium tumefaciens* AGL1 strain by electroporation using the MicroPulser Electroporator (www.bio-rad.com) according to the manufacturer’s instructions. The AGL1-pCS-GFP and AGL1-pCS-hpGF+GFP were used to transform FsK conidia as previously described (Zhang *et al*., 2015).

### *In vitro* transcription of sGFP dsRNA

For the generation of the in vitro transcribed sGFP dsRNA, genomic DNA was extracted from FsK-sGFP (Sesma and Osbourn, 2004) and used as template for PCR with KAPA Taq DNA Polymerase (www.sigmaaldrich.com) with the T7 promoter-containing primers 5’-taa tac gac tca cta tag gga gaC GTA AAC GGC CAC AAG TTC AGC-3’ and 5’-taa tac gac tca cta tag gga gaG TGG CGG ATC TTG AAG TTC ACC-3’ (T7 promoter sequence with lowercase). The T7 promoter-containing 491 bp amplicon was then used as template in the MEGAscript™ RNAi Kit (www.thermofisher.com) for the generation of a 445 bp sGFP dsRNA.

### In vitro RNAi assay

In 24 wells of a 96-well plate, FsK-sGFP conidia were added (in each well, 6 conidia diluted in 100 μl PDB/100). In 12 wells containing these FsK-sGFP conidia, in vitro transcribed sGFP dsRNA was added (100 μl, 1 ng/μl) (dsRNA application samples). In the remaining 12 wells containing FsK-sGFP conidia, 100 μl water was added (control samples). The 96 well was covered with a removable membrane and incubated at 28° C. At timepoints 0-24-48 hpa, the plate was subjected to fluorometric analysis using the using the Varioskan™ LUX multimode microplate reader (www.thermofisher.com).

### Bisulfite sequencing

Genomic DNA from the fungus (20 ng) or the plant (100 ng) was used for bisulfite sequencing analysis using the EZ DNA Methylation-Gold Kit (www.zymoresearch.com) according to the manufacturer’s instructions and as previously described (Dalakouras *et al*., 2016). Essentially, for the cis-RdDM bisulfite analysis on FsK, the primers 5’-AAT CTC CAR TRR RTA CAC TAT TC-3’ and 5’-CCT CCT TRA AAT CRA TTC CCT TAA-3’ were used, whereas for the trans-RdDM bisulfite analysis on FsK and Nb-16C the primers 5’-AGT GGA GAG GGT GAA GGT GAT G-3’ and 5’-CCT CCT TRA AAT CRA TTC CCT TAA-3’ were used in a PCR reaction with ZymoTaq PreMix (www.zymoresearch.com) according to the manufacturer’s instructions. The occuring 262 bp and 311 bp amplicons for cis-RdDM and trans-RdDM, respectively, were cloned into pGEM®-T Easy Vector (worldwide.promega.com) and for each analysis 5-10 clones were subjected to Sanger sequencing.

### Small RNA sequencing

Sequencing of small RNAs from fungal RNA (small RNA fraction) was performed by GenXPro (https://genxpro.net/) as previously described (Dalakouras *et al*., 2016).

## Results and Discussion

### FsK colonizes the root system of Nicotiana benthamiana and stimulates plant growth

During the colonization process of its host plants, the fungus penetrates the root and grows in the root cortex and proliferates even in the vascular system of root system (Skiada *et al*., 2019). In legumes, efficient colonization by FsK is dependent on the common symbiotic signalling pathway (Skiada *et al*., 2020), typically used by rhizobia and arbuscular mycorrhizal fungi. Notably, although not yet explained for, fungal growth in tomato is restricted to the root system and extends only to the crown and not to the stem and leaf tissues (Kavroulakis *et al*., 2007). Here, we investigated the capacity of FsK to colonize another member of the Solanaceae, *Nicotiana benthamiana*, which is a widely used model plant for RNAi studies (Philips *et al*., 2017). Similar to tomato, upon root-inoculation, the fungal endophyte colonized the root system but failed to expand to the shoot system (Figs 1a, 1b and S1). Interestingly, the FsK-colonized plants exhibited considerably stimulated growth, at least up to 4 weeks post inoculation (wpi) when grown in both non-sterile compost (Fig. 1c) and sterile sand (Fig. S2), underpinning the beneficial effect of FsK to this host, at least in terms of biomass production.

**Fig. 1.**
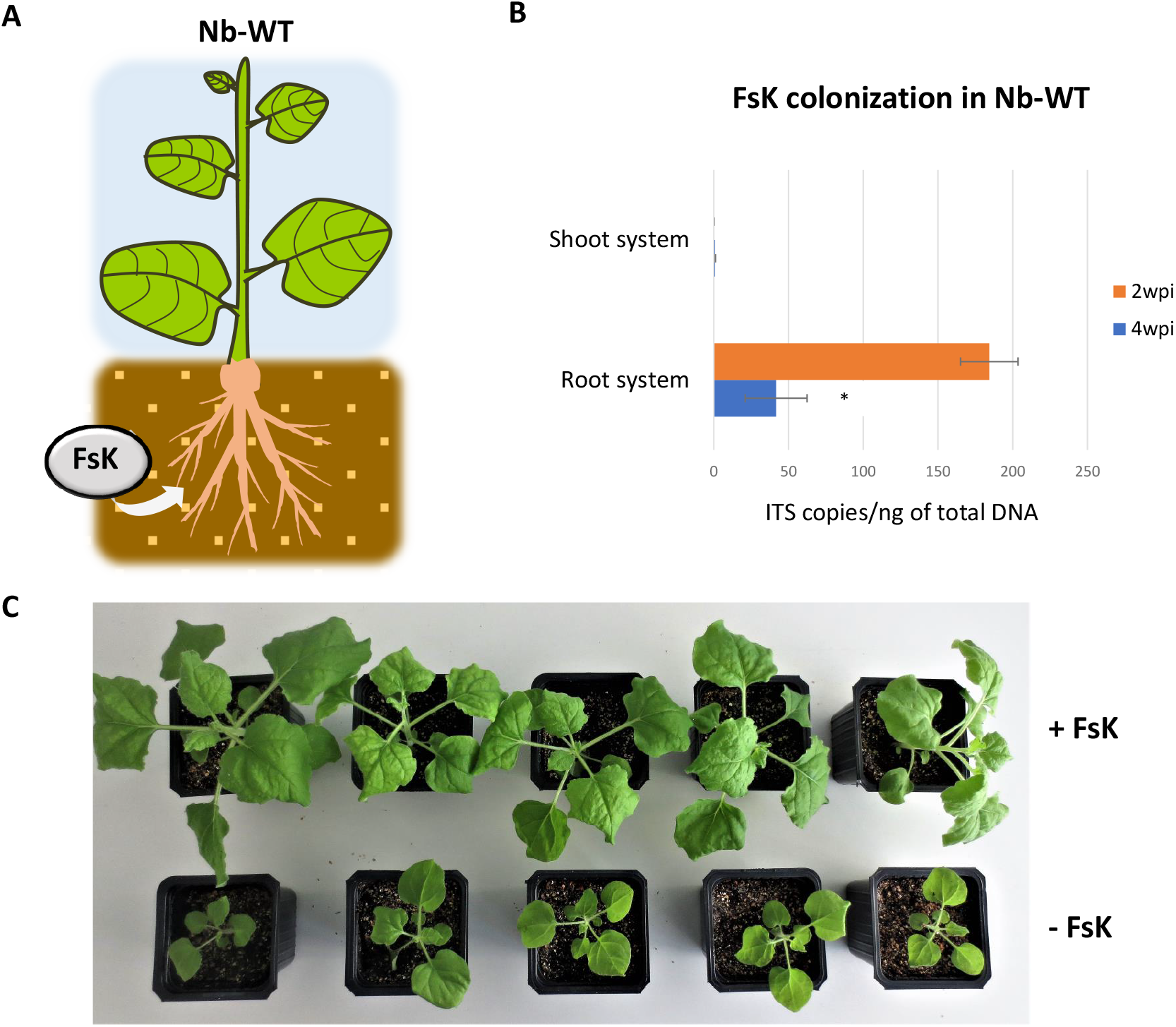
Colonization of Nb-WT by FsK. (A) Schematic representation of the colonization assay. (B) Quantification of fungal colonization in shoot and root system at 2 and 4 wpi. (C) Impact of the FsK in growth of Nb-WT 4 wpi.

### FsK encodes the core RNAi components

Despite being largely conserved among eukaryotes, not all fungi encode the core RNAi pathway; indeed, *Saccharomyces cerevisiae* lacks DCLs, AGOs and RDRs (Drinnenberg *et al*., 2009). *Ustilago maydis* also lacks DCLs, AGOs and RDRs, in contrast, surprisingly, to its close relative *U. hordei* (Laurie *et al*., 2008). Furthermore, miRNAs have been identified in most fungal species but not in the basal fungus *Mucor circinelloides* (Torres-Martinez and Ruiz-Vazquez, 2017). Interestingly though, whereas RNAi-deficient mutants of most ascomycetes and basiodiomycetes are not impaired in vegetative growth and development, sexual differentiation and response to stress, *M. circinelloides* is (Ruiz-Vazquez *et al*., 2015). These being said, the mechanistic details and role of RNAi in fungal kingdom can be unusually diverse.

To examine whether FsK encodes the core RNAi machinery, we performed transcriptome-validated genome annotation (BioProject PRJNA796177, Tsiouri and Papadopoulou, unpublished results) and identified two DCLs (FsKDCL1 and FsKDCL2), two AGOs (FsKAGO1 and FsKAGO2) and four RDRs (FsKRDR1-4) (Fig. 2a). FsKDCL1 and FsKDCL2 contain the Dicer-like protein structures with a Dead-like helicases superfamily domain box (DEXDc) box, a helicase superfamily c-terminal domain (HELICc), and two ribonuclease III domains (RIBOc) responsible for the cleavage of dsRNA precursors into sRNAs (Paturi and Deshmukh, 2021). Both FsKAGO1 and FsKAGO2 proteins contain PAZ and PIWI domains; PAZ recognizes the 3’ end of sRNAs while PIWI exhibits an RNaseH-like endonucleolytic activity and mediates target cleavage (Wu *et al*., 2020). All four FsKRDRs contain the RdRP/RDR domain, which is highly conserved in fungi (Chen *et al*., 2015). FsKRDR2 and FsKRDR3 contain the DLDGD motif, which is often encountered in plants, whereas FsKRDR1 and FsKRDR4 contain the DYDGD motif, which is more common in fungi (Wassenegger and Krczal, 2006). To explore the molecular evolution of these proteins, we performed phylogenetic analyses of DCL, AGO and RDR proteins including *Fusarium graminearum* and *Neurospora crassa* (Fig. 2b). Our analysis showed that FsKDCL1 is related to FgDCL1 and NcDCL1 that function in the meiotic silencing by unpaired DNA (MSUD) pathway (Fig. 2b) (Alexander *et al*., 2008), whereas FsKDCL2 is closer to FgDCL2 and NcDCL2 which have a prominent role in RNAi by processing of dsRNAs into siRNAs (Chen *et al*., 2015). FsKAGO1 is closely related to FgAGO1 and NcQDE2 that are loaded with dsRNA-processed siRNAs during RNAi, whereas FsKAGO2 is closer to FgAGO2 and the *N. crassa* SMS2 that are involved in MSUD (Lee *et al*., 2003). Of note, FsKRDR4 is closely related to NcQDE1 which is essential for quelling and suggested to be functionally related to plant RDR6 (Wassenegger and Krczal, 2006).

**Fig. 2.**
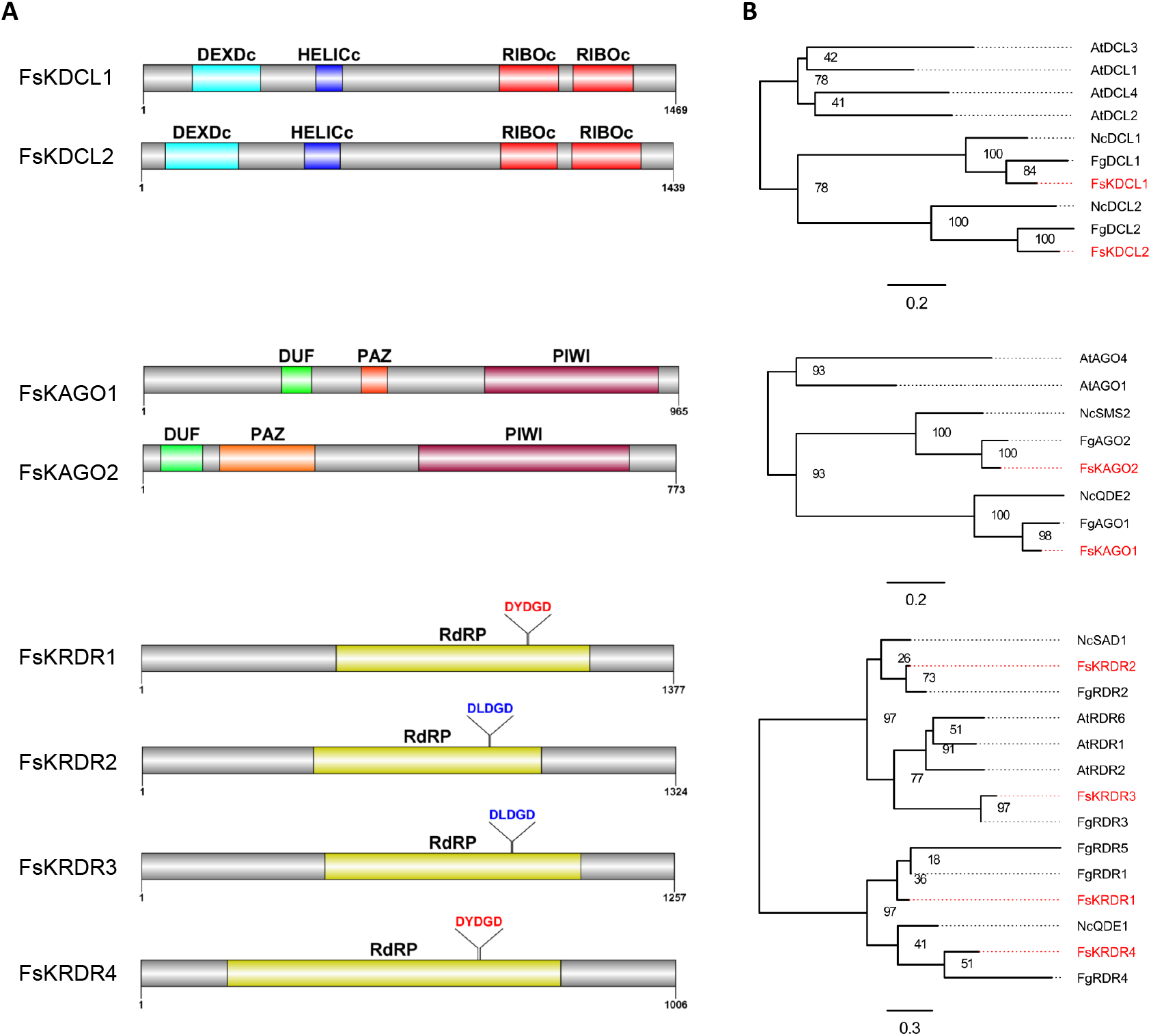
Identification of FsK RNAi core machinery. (A) Schematic representation of FsK DCL, AGO and RDR proteins using DOG1.0 software (Ren *et al*., 2009). (B) Maximum likelihood phylogenies of the FsK (indicated red), *Fusarium graminearum, Neurospora crassa* and *Arabidopsis thaliana* (as an outgroup member) DCL, AGO and RDR proteins using the LG model matrix and 100 bootstrap replicates for assessing branch support.

### FsK takes up RNAi molecules from its environment

In order to test the functionality of FsK’s RNAi machinery, an *in vitro*-transcribed 445 bp sGFP dsRNA was applied to a sGFP-expressing FsK, sGFP being a GFP variant that contains a serine-to-threonine substitution at amino acid 65, optimized for use in fungi (Sesma and Osbourn, 2004) (Figs 3a, 3b). Fluorometric analysis revealed that the sGFP expression levels dropped to almost 50% 24 hours post application (hpa) (Fig. 3c). These data suggested that the externally applied dsRNA was processed by fungal DCLs into siRNAs that were loaded onto fungal AGOs to mediate cleavage of the sGFP mRNA. However, no further decrease of sGFP levels could be observed at later timepoints (48 hpa), reminiscent of similar observations in *F. asiaticum* (Song *et al*., 2018) and implying the absence an active RDR-mediated self-reinforcing mechanism of RNAi that could ensure ongoing RNAi even at the absence/degradation of the initial dsRNA input.

**Fig. 3.**
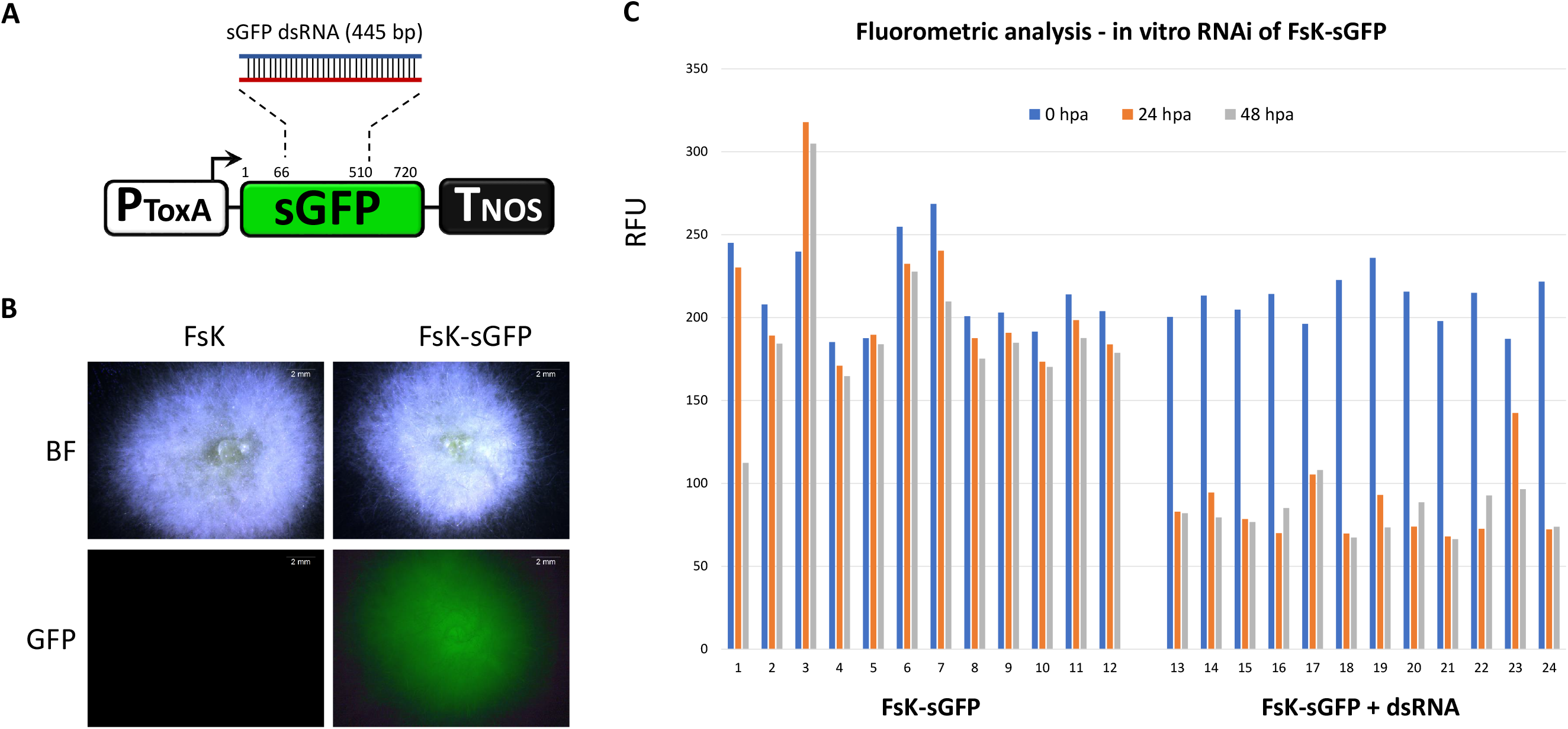
In vitro RNAi in FsK-sGFP. (A) Schematic representation of the sGFP transgene that is present in FsK-sGFP. PToxA: promoter for the promoter from *Pyrenophora tritici-repentis* ToxA gene; sGFP: GFP variant that contains a serine-to-threonine substitution at amino acid 65; TNOS: terminator for the nopaline synthase gene. The 445 bp fragment chosen for in vitro transcription of dsRNA is depicted. (B) Stereoscopic observation of sGFP fluorescence. (C) Fluorometeric analysis for in vitro RNAi in FsK-sGFP. Vertical axis: RFU: relative fluorescence unit, calculated as the ratio of sGFP-indicative fluorescence (excitation 488 nm, emission 515 nm) to growth-indicative absorbance (wavelength 595 nm). Horizontal axis: 1-12: 12 wells containing FsK-sGFP conidia. 13-24: 12 wells containing FsK-sGFP conidia plus 100 ng (each well) sGFP dsRNA.

Overall, these data suggest not only that the RNAi machinery in FsK is functional but also that FsK is able to take up RNAi molecules from its environment. Not all fungi are able to take up RNA molecules from their environment; *Colletrotrichum gloesporiodes, Trichoderma virens* and *Phytophtora infestans* being some notable examples that fail to do so (Qiao *et al*., 2021). Of note, fungi that are indeed able to receive RNAi molecules from their environment are not only promising candidates for RNAi-based fungicidal control (Šečić and Kogel, 2021) but also likely partners in an RNAi-based cross-kingdom communication with their host (He *et al*., 2021).

### FsK processes hairpin RNA transcripts into siRNAs that trigger mRNA degradation but not DNA methylation in the fungal hyphae

In order to examine the mode of dsRNA processing in the endophyte, FsK was transformed with a transgene comprised of a full length green fluorescent protein (GFP) corresponding to mGFP5-ER (Haseloff and Siemering, 2006) and a hairpin (hp) construct of the first 332 bp of GFP (hpGF) (Fig. 4a). In this setup, the hpGF locus served as the RNAi-trigger while the GFP locus as the RNAi-target. Small RNA sequencing (sRNA-seq) in three independent FsK-hpGF+GFP transformants (#6, #7, #27) revealed the accumulation of GF siRNAs (perfectly matching the GF region) having variable sizes from 18-30 nt but predominantly of 21-nt, 22-nt and 24-nt (Figs 4b, 4c and S3). This finding was reminiscent of the situation in plants, where hpRNAs are typically processed by DCLs to 21-, 22- and 24-nt siRNAs (Fusaro *et al*., 2006). To the best of our knowledge, similar sRNA-seq studies in fungi, aiming to reveal the mode of processing of a specific hpRNA/dsRNA, are absent; yet, genome-wide sRNA-seq studies (identifying siRNAs, miRNAs but also DCL-independent sRNAs) reveal a remarkably diverse pattern, with prominent size classes ranging from 19-22-nt in *Penicillium chrysogenum* (Dahlmann and Kück, 2015), to 22-25-nt in *S. pombe* (Djupedal *et al*., 2009) and 27-28-nt in *F. graminearum* (Chen *et al*., 2015). Our analysis does not allow us to identify whether all sRNA size classes are actual DCL products (e.g., they could represent degradation products) or whether they all exhibit biological activity. Yet, it is reasonable to assume that FsKDCL2 generated the bulk of sRNAs (Chen *et al*., 2015), of which the most prominent size class (21-nt) seems to undertake the major burden for RNAi activity.

**Fig. 4.**
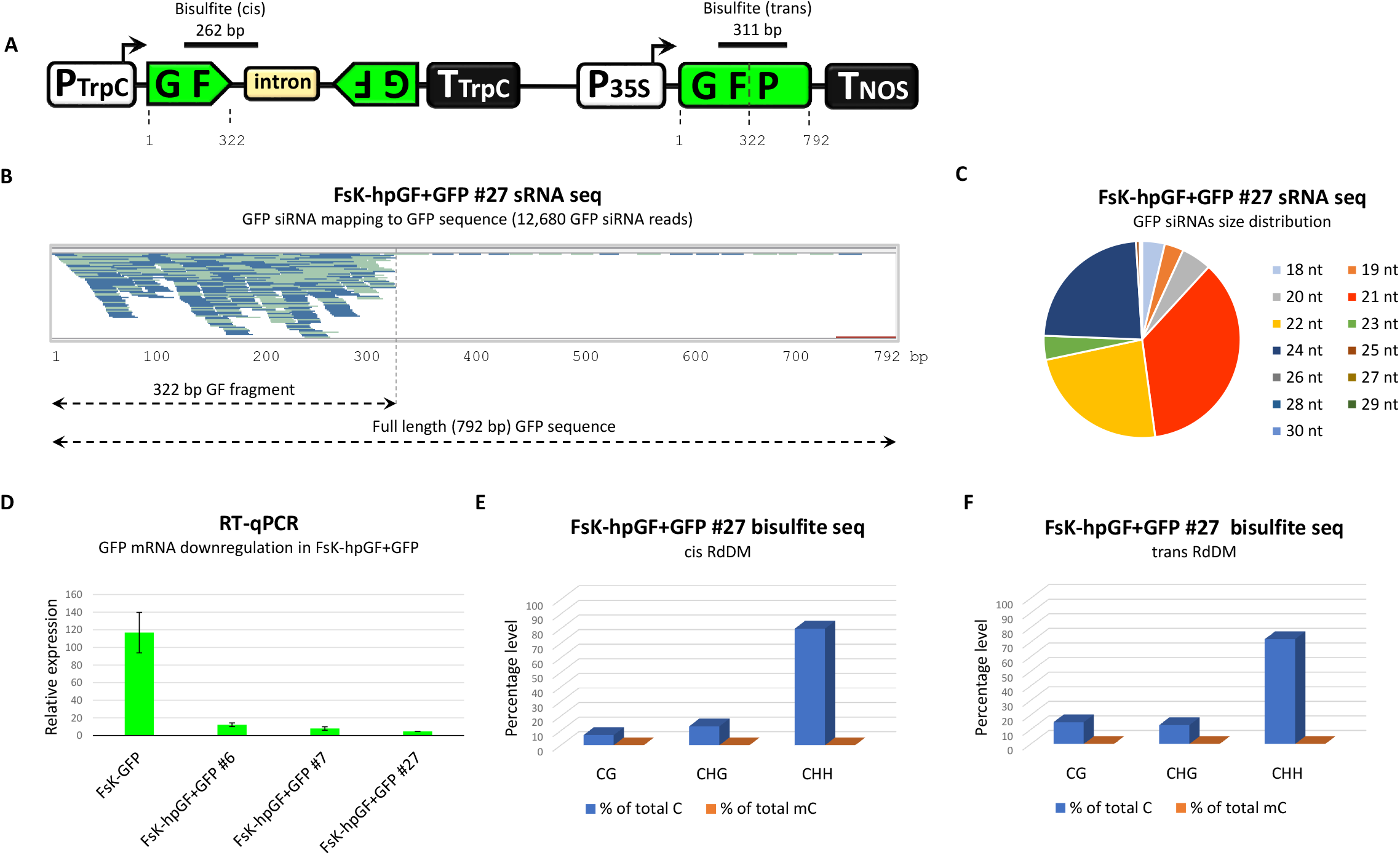
Characterization of FsK RNAi machinery. (A) Schematic representation of the hpGF+GFP transgene. PTrpC: promoter for the Promoter for *Aspergillus nidulans* trpC gene; GF: 322 bp fragment of the GFP; intron: *Magnaporthe grisea* cutinase gene intron; TTrpC: promoter for the Promoter for Aspergillus nidulans trpC gene, P35S: Cauliflower mosaic virus 35S promoter; GFP: full-length (792 bp) green fluorescent protein (mGFP-ER version); TNOS: terminator for the nopaline synthase gene. FsK-hpGF+GFP transformants contain the full length hpGF+GFP transgene, whereas FsK-GFP transformants contain only the P35S-GFP-TNOS part of the transgene. (B) SRNA-seq in FsK-hpGF+GFP #27. All sRNA reads of 18-30-nt fully matching to GFP region are depicted. With light blue the siRNA reads in plus polarity, with dark blue the siRNA reads in minus polarity. The Tablet software (Milne *et al*., 2013) was used for visualization of the sRNA reads. (C) Pie graph of the 18-30 nt GFP siRNAs in FsK-hpGF+GFP #27. (D) RT-qPCR for the estimation of GFP mRNA downregulation in FsK-hpGF+GFP compared to FsK-GFP. (E) Bisulfite sequencing for cis RdDM. (F) Bisulfite sequencing for trans RdDM.

To evaluate this RNAi activity, we measured the GF siRNA-mediated downregulation of GFP mRNA in three independent FsK-hpGF+GFP transformants (#6, #7, #27) when compared to FsK-GFP (transformed with a cassette lacking the hpGF transgene) (Fig. 4a). Indeed, GFP expression was virtually eliminated in all FsK-hpGF+GFP transformants (Figure 3D). Of note, we detected GF siRNAs but no or negligible P siRNAs that could had potentially emerged upon the FsKRDR processing on the GF siRNA-targeted GFP transcript (Figure 4b). This is in contrast to the situation in plants (de Felippes and Waterhouse, 2020) but in agreement with similar reports in *F. asiaticum* (Song *et al*., 2018), suggesting the absence of an active RDR-based mechanism in FsK.

Typically, the onset of RNAi and the accumulation of siRNAs leads to RdDM in plants (Dalakouras and Vlachostergios, 2021). DNA methylation also occurs in some, but not all, fungi, and usually in repetitive sequences (Bewick *et al*., 2019). Yet, such DNA methylation is considered to be dispensable of RNAi molecules, thus fungi have been considered to lack a *bona fide* RdDM mechanism (Nai *et al*., 2020). Nevertheless, recent advances challenge this assumption; indeed, sRNA-dependent RdDM-like phenomena has been detected, at least in *Pleurotus tuoliensis and P. eryngii var. eryngii* (Basiodiomycetes) (Zhang *et al*., 2018) and *Puccinia graminis* (Ascomycetes) (Sperschneider *et al*., 2021). Accordingly, and given the abundant accumulation of GF siRNAs in FsK-hpGF+GFP, we were interested to see whether they could trigger RdDM of cognate DNA sequences. To analyze cis-RdDM (at the locus generating the siRNAs), we chose a 262 bp fragment of the hpGF transgene (Fig. 4a). For trans-RdDM (at a locus that does not generate siRNAs but is homologous to them), we chose a 311 bp fragment of the GFP transgene (Fig. 4a). Whereas CG and CHG methylation can be maintained in an RNAi-independent manner (Law and Jacobsen, 2010), CHH methylation is the hallmark of ongoing de novo RdDM (Pelissier *et al*., 1999), and both cis and trans fragments under analysis were rich in asymmetric CHH context (80% for cis and 72% for trans) (Figs 4e, 4f). However, bisulfite sequencing revealed the absence of methylated cytosines in any sequence context (CG, CHG, CHH), at neither cis (Fig. 4e) nor trans (Fig. 4f) loci, suggesting that no RdDM takes place in FsK, at least in our experimental setup. It has been suggested that fungal proteins with de novo methyltransferase (DNMT) and/or helicase-like Snf2 family domains may be involved in RdDM-like pathways in fungi (Nai *et al*., 2020). However, were unable to detect such genes in the FsK genome, underpinning the conclusions obtained from bisulfite sequencing about the absence of an active RdDM mechanism in FsK.

### FsK translocates RNAi signals to its host to induce systemic RNAi and epigenetic changes of a reporter gene

Establishment of mutualistic associations between fungi and their host requires genetic and epigenetic reprogramming as well as metabolome modulation of both by the exchange of effector molecules (Kloppholz *et al*., 2011). Indeed, RdDM is essential in Arabidopsis to establish a beneficial relationship with the root-colonizing *Trichoderma atroviride* while DNA methylation and histone modifications are required for plant priming by the beneficial fungus against *B. cinerea* (Rebolledo-Prudencio *et al*., 2021). Importantly, it was just recently shown that during the mutualistic interaction of the ectomycorrhizal fungus *Pisolithus microcarpus* with *Eucalyptus grandis*, a fungal miRNA, Pmic_miR-8, targets the host NB-ARC domain containing transcripts in a cross-kingdom RNAi manner (Wong-Bajracharya *et al*., 2022). Reminiscent of this, an in silico study predicted that the beneficial arbuscular mycorrhizal fungi *Rhizophagus irregularis* produces sRNAs that have 237 candidate targets in the host plant *Medicago truncatula*, including specific mRNAs known to be modulated in roots upon AMF colonization (Silvestri *et al*., 2019). Similarly, a recent study based on transcriptome and sRNA profile change analysis during the onset of the mutualistic interaction between the beneficial root endophyte *Serendipita indica* with its host *Brachypodium distachyon*, suggested that interaction-induced sRNAs in both organisms may underlie reciprocal targeting of genes related to plant development and fungal growth and nutrient acquisition (Secic *et al*., 2021). Thus, it is very likely that, similar to fungal pathogens (Cai *et al*., 2019), beneficial fungal endophytes also display an RNA-based communication with their hosts. However, clear evidence of actual RNAi molecule translocation and concomitant cross-kingdom RNAi between a beneficial fungal endophyte and its host has been lacking to this date.

In order to address this question, we resorted to the GFP-expressing *N. benthamiana* plant line 16C (Nb-GFP) (Voinnet and Baulcombe, 1997), as an RNAi sensor system. Nb-GFP carries a 35S-driven mGF5-ER transgene (Fig. 5a) and is a well-studied RNAi model plant that allows the monitoring of systemic RNAi (i.e. spreading of RNAi to tissues other than those where RNAi initially occurred) by observation of the presence or abolishment of GFP expression under ultraviolet light. When Nb-GFP plants were inoculated with FsK-hpGF+GFP (Fig. 5a), we could record the following outcomes: (i) no visible RNAi (45% of the plants, 6 wpi), (ii) spot-like RNAi (45% of the plants, 4 wpi), (iii) vein-restricted RNAi (5% of the plants, 4 wpi) and (iv) full-tissue RNAi (5% of the plants, 4 wpi) (Fig. 5b). Colonization of Nb-GFP plants with non-transformed FsK and/or FsK-sGFP failed to trigger any visible RNAi phenotype even after 10 wpi, suggesting that not the mere presence of the endophyte but the RNAi molecules it expresses are responsible for the induction of RNAi phenotypes in its host.

**Fig. 5.**
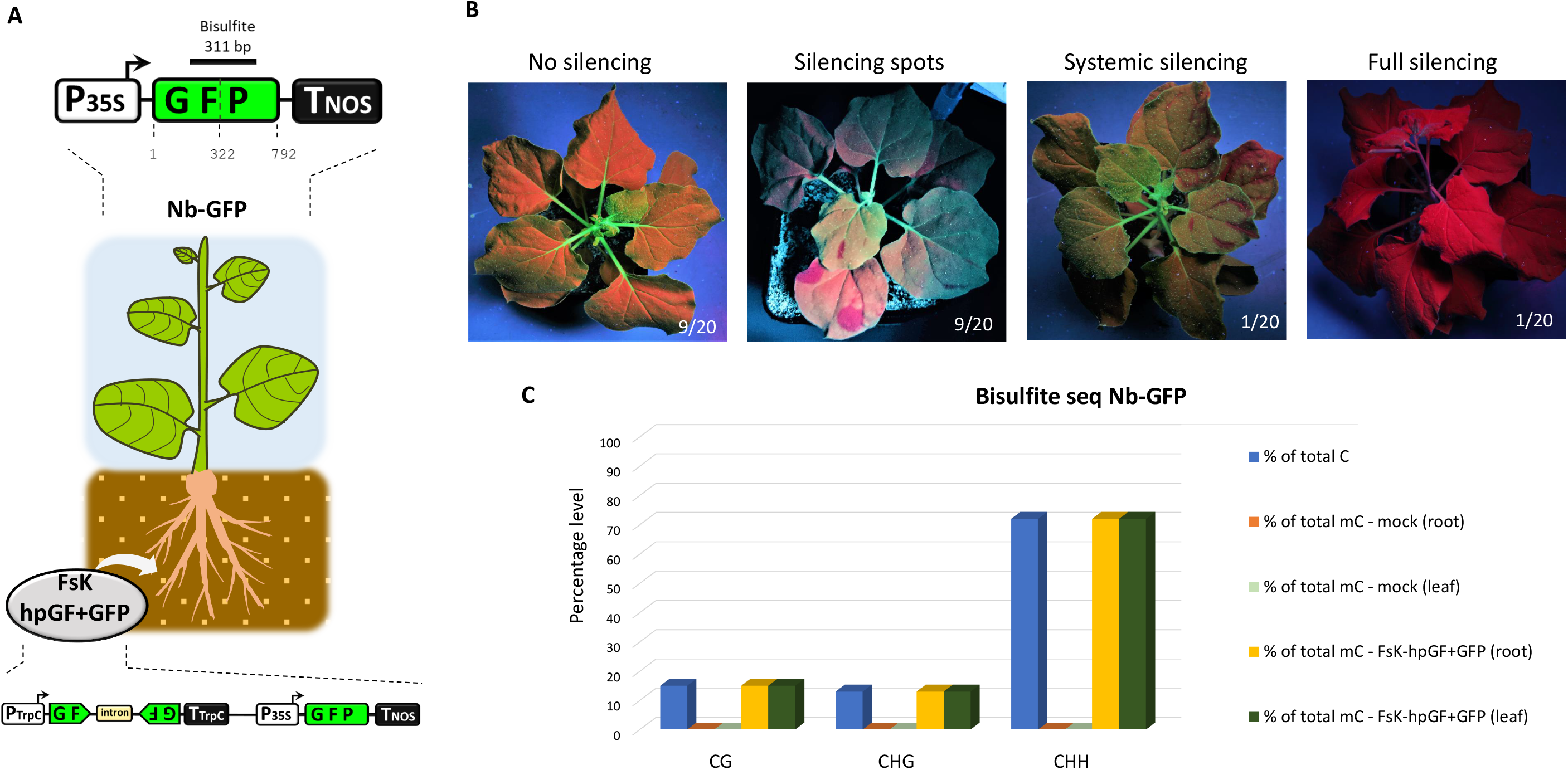
FsK-hpGF+GFP colonization of Nb-GFP. (A) Schematic overview of the colonization assay. (B) Systemic silencing phenotypes under ultraviolet light 4-6 wpi. (C) Bisulfite sequencing in the host GFP transgene in both roots and leaves in silenced Nb-GFP plants.

RNAi in plants is tightly coupled to RdDM (Dalakouras and Vlachostergios, 2021; Jones *et al*., 1999). Accordingly, bisulfite sequencing analysis of leaf and root tissues from the fully silenced Nb-GFP plants disclosed the dense (100%) onset of DNA methylation in the GFP region (Fig. 5a) in every sequence context: CG, CHG and CHH (Fig. 5c). Overall, these data clearly show that the endophyte triggered not only mRNA degradation but also DNA methylation of a host reporter gene. We favor the scenario that FsK-hpGF+GFP translocated RNAi signals (dsRNAs but most likely siRNAs) to the roots of Nb-GFP initiating local RNAi of the host GFP. Importantly, once present in the plant cells and upon targeting the host GFP transcript for silencing, these endophyte-derived primary siRNAs culminated in the generation of host-derived RDR-mediated secondary siRNAs (as implied by the RdDM pattern, see below). Whether the recorded RdDM in the root tissues was induced by the endophyte-derived primary or the host-derived secondary siRNAs is not clear. Yet, the fact that RdDM could be detected not only in the GF but also in the P region (Figure 5a, 311 bp bisulfite fragment covering both GF and P regions) strongly implies in favor of transitive host-derived secondary siRNAs imposing RdDM. Now, siRNAs are mobile moieties; they can move cell-to-cell through the plasmodesmata and through the vasculature to distant parts of the plant (Voinnet, 2022; Voinnet and Baulcombe, 1997). The establishment of systemic silencing in the upper parts of the plant (which FsK fails to colonize, Fig. 1) suggests that mobile siRNA signals from the root entered the phloem to reach shoot tissues. Most likely both endophyte-derived primary siRNAs and host-derived secondary siRNAs could have played the role of the mobile systemic signal (Devers *et al*., 2020). However, it is unlikely that the mere presence of endophyte-derived primary siRNAs alone could trigger systemic silencing; it rather seems that a certain quantitative siRNA threshold needs to be surpassed for the onset of systemic silencing (Kalantidis *et al*., 2006), rendering the abundant presence of host-derived secondary siRNAs indispensable. Importantly, establishment of systemic RNA in the receiving tissues requires RDR6 (Schwach *et al*., 2005). Thus, in the receiving tissues, the primary/secondary siRNAs triggered a RDR6-mediated generation of (host-derived) tertiary siRNAs, ensuring the efficient establishment of GFP mRNA degradation and DNA methylation.

### Conclusion

Here, we have characterized the RNAi core machinery of a fungal endophyte and we provide solid evidence that it translocates RNAi signals to its host to trigger systemic silencing and epigenetic modifications. To prove the concept, we have employed an artificial RNAi sensor system; future studies coupling sRNAome, degradome and methylome analysis will be required to pinpoint the nature of the endogenous fungal sRNAs (siRNAs and/or miRNAs) that are translocated to the host, which host genes are targeted for transcriptional and/or post-transcriptional silencing and how this process is ultimately translated into a beneficial phenotype. Our data may well reflect a so far unrecognized pathway according to which endophytes establish the symbiosis and/or impose their beneficial impact by translocating RNA molecules that modulate host gene expression and affect the epigenome’s plasticity. RNAi-mediated communication between plants and their interacting organisms is much more widespread than previously thought and may account for the improved plant performance often observed in the presence of certain associated microbiota.

## Supplementary Data

The following supplementary data are available at JXB online.

Fig. S1. Colonization of FsK-sGFP in Nb-WT and stereoscopic observation of sGFP fluorescence in various tissues

Fig. S2. Impact of FsK colonization of Nb-WT plants grown in sterile sand in magenta boxes.

Fig. S3. Small RNA sequencing in three FsK-hpGF+GFP transformants. (a) Mapping of sRNAs in GFP. (b) Size distribution of GFP sRNAs.

Fig. S4. Systemic silencing phenotypes upon colonization of FsK-GF+GFP in Nb-GFP plants 4-6 wpi.

## Acknowledgments

This work was supported by funds from the EU Horizon 2020 Marie Skłodowska-Curie fellowship (RNASTIP, Grant ID 793186) and the EU Horizon 2020 PRIMA program (INTOMED, Grant ID 1534).

## Author Contributions

A.D. and K.K.P. designed research; A.D., A.K., M.A., E.D.., A.M. M.G. and E.D., performed research; A.D., O.T., S.V. and KKP analyzed data; A.D. and K.P.P. wrote the paper. All authors reviewed and approved the manuscript.

## Data availability

All sequencing data supporting the findings of this study are deposited to Zenodo (https://doi.org/10.5281/zenodo.6088855)

---

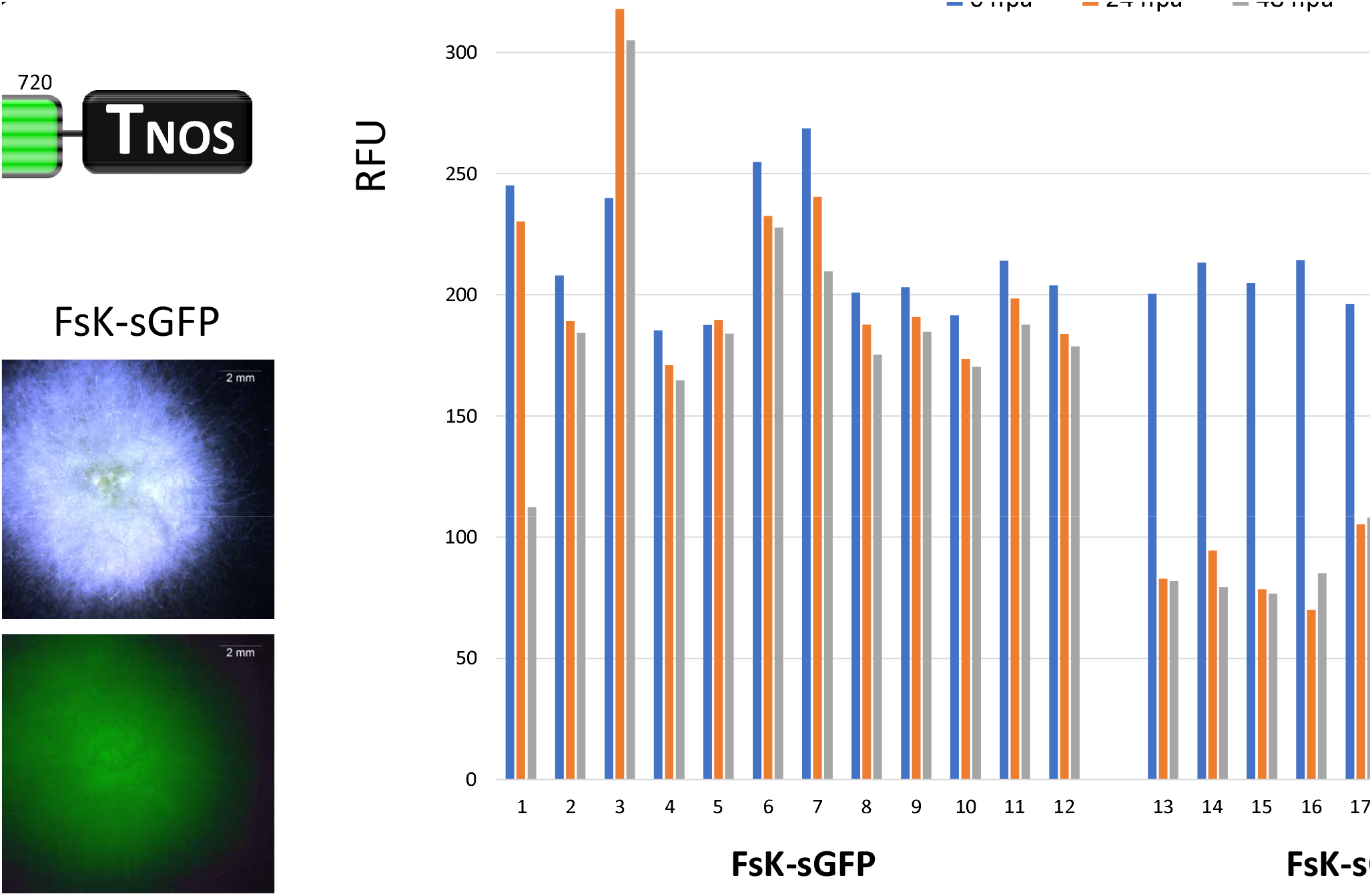

