## Supplementary material for "Systemic silencing and DNA methylation of a host reporter gene induced by a beneficial fungal root endophyte": Suppl. Fig 1

Fig. S1

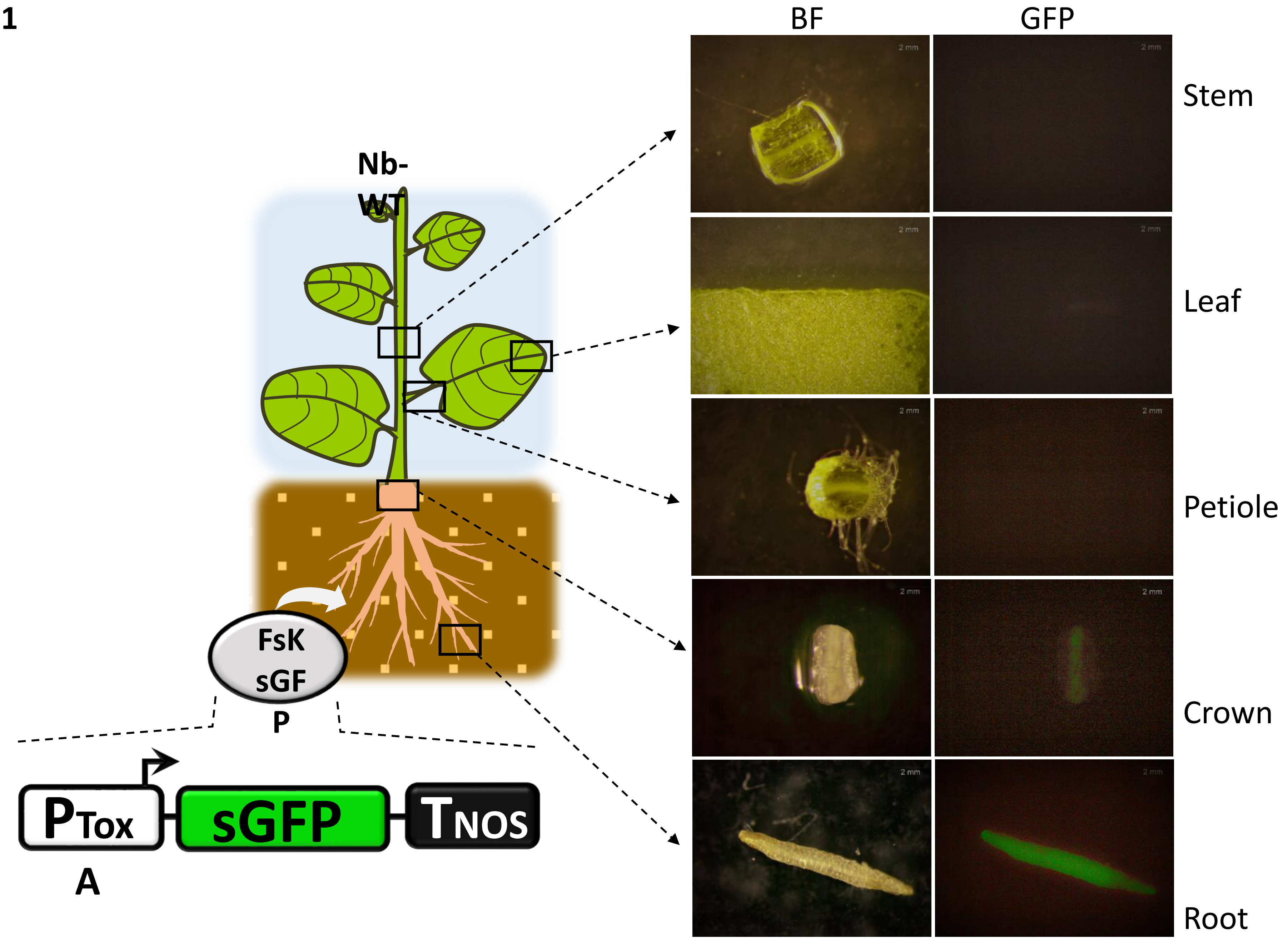


Fig. S1. Colonization of FsK-sGFP in Nb-WT and stereoscopic observation of sGFP fluorescence in various tissues
