## Supplementary figures and images for "Systemic silencing and DNA methylation of a host reporter gene induced by a beneficial fungal root endophyte"

### Suppl. Fig 2

**Fig. S2**

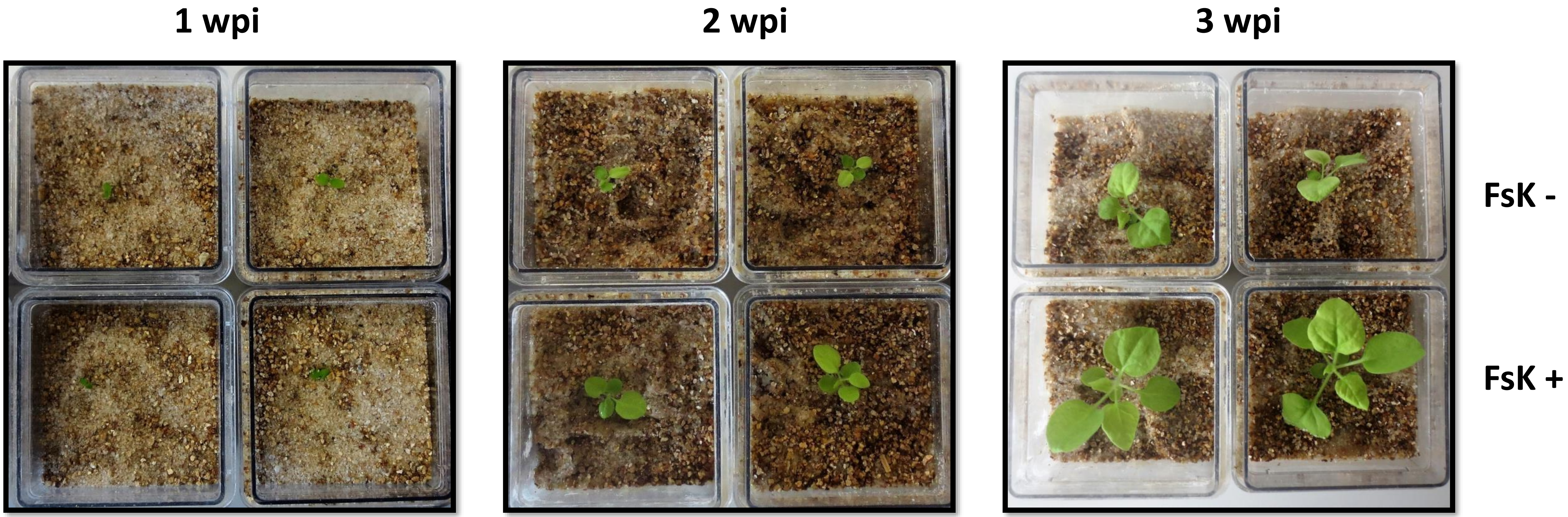


Fig. S2. Impact of FsK colonization of Nb-WT plants grown in sterile sand in magenta boxes

### Suppl. Fig 4

**Fig. S4**

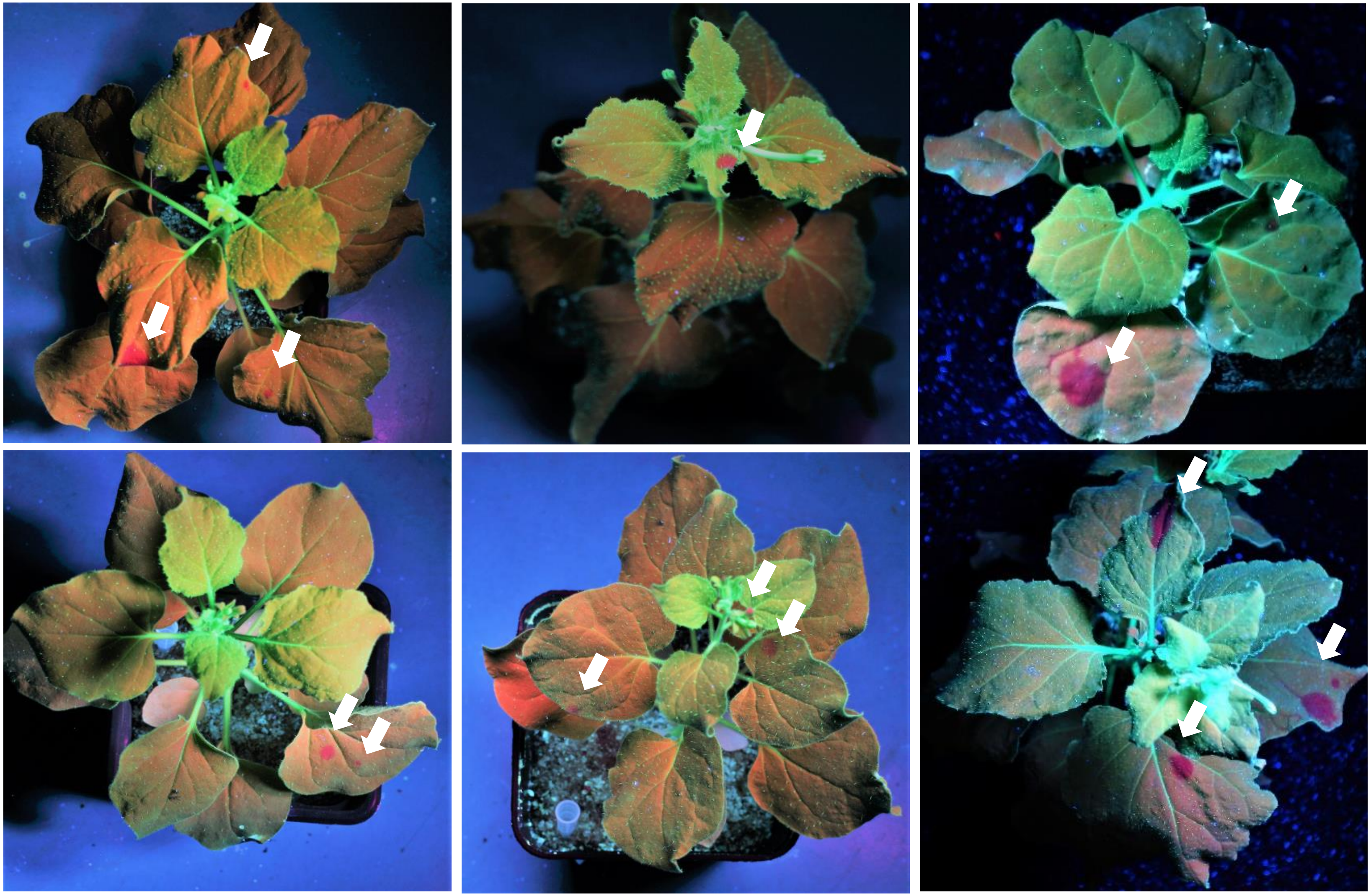

Fig. S4. Systemic silencing phenotypes upon colonization of FsK-GF+GFP in Nb-GFP plants 4-6 wpi
