## Supplementary material for "Systemic silencing and DNA methylation of a host reporter gene induced by a beneficial fungal root endophyte": Suppl. Fig 3

Fig. S3

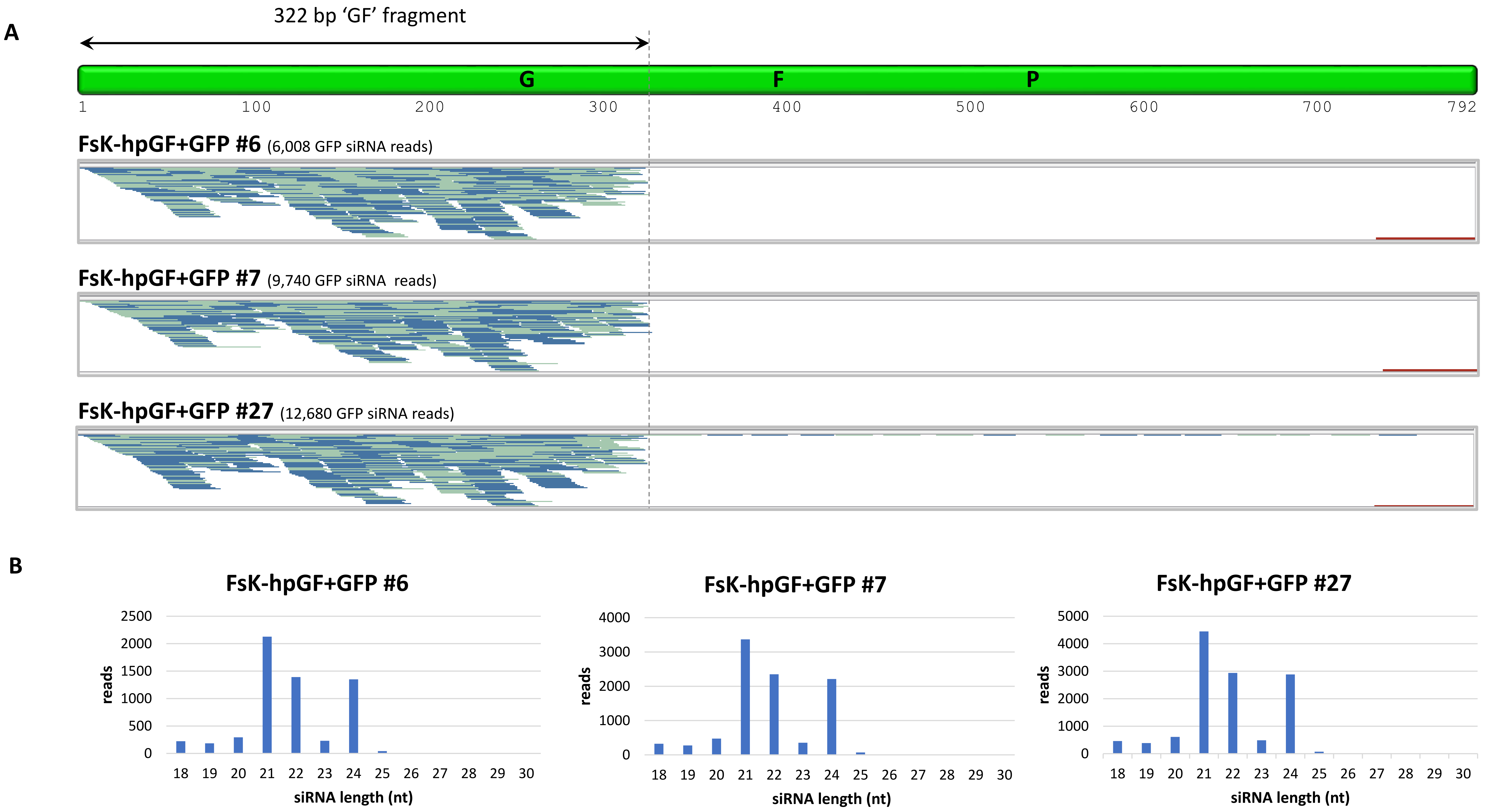


Fig. S3. Small RNA sequencing in three FsK-hpGF+GFP transformants. (a) Mapping of sRNAs in GFP. (b) Size distribution of GFP sRNAs
